# GATSBI: Improving context-aware protein embeddings through biologically motivated data splits

**DOI:** 10.64898/2026.02.13.705830

**Authors:** Gowri Nayar, Russ B. Altman

**Affiliations:** Department of Biomedical Data Science, Stanford University, CA, USA; Department of Bioengineering, Stanford University, CA, USA; Department of Genetics, Stanford University, CA, USA; Department of Medicine, Stanford University, CA, USA

**Keywords:** Heterogeneous protein networks, Graph attention networks, Context-aware protein embeddings

## Abstract

**Motivation:** Understanding protein function requires integrating diverse biological evidence while accounting for strong contextual dependence. Recent protein embedding methods increasingly leverage heterogeneous biological networks, yet their evaluation protocols often fail to reflect the specific biological tasks for which the embeddings are intended. Prediction of missing interactions, annotation of new proteins, and discovery of functional modules require fundamentally different data partitions, such as edge-masked versus node-held-out splits. Moreover, most approaches report performance primarily on well-studied proteins, where computational predictions are least needed, risking substantial overestimation of real-world utility.

**Results:** We introduce a graph attention–based framework (Gatsbi) to construct context-aware protein embeddings from integrated protein–protein interactions, co-expression, sequence representations, and tissue-specific associations. Using task-aligned evaluation protocols, we show that models trained with biologically appropriate partitions achieve markedly better generalization. Across interaction, function, and functional set prediction, Gatsbi consistently outperforms existing pretrained embeddings for both well-studied and understudied proteins, with the largest gains observed for the understudied regime and under inductive node-held-out evaluation. To enable broad reuse, we provide the learned embeddings for download for application to other protein prediction tasks.

**Availability and Implementation:** https://github.com/Helix-Research-Lab/GATSBI-embedding

## 1. Introduction

Protein function depends on cellular and biological context [1]. To capture this complexity, computational models increasingly integrate heterogeneous data—protein–protein interactions, co-expression, transcriptomics, and sequence or structural features—to learn context-aware protein embeddings [2]. These embeddings place functionally related proteins near one another in representation space, enabling downstream tasks such as interaction prediction, functional annotation, and disease association. This is especially important for understudied proteins, which lack direct experimental characterization and rely on information transfer from well-studied neighbors [3]. Because most future predictions concern such proteins, performance in this regime is a key measure of real-world utility.

Biological evidence is inherently relational, making heterogeneous networks a natural way to integrate diverse data sources such as physical interactions, co-expression, and tissue-specific associations [4]. These networks preserve both the diversity of biological evidence and the structure of functional relationships between proteins. Graph attention networks (GATs) are well suited to this setting: they update each proteins representation by aggregating information from its neighbors with learned attention weights that reflect the importance of each biological relationship [5]. By stacking multiple attention layers, GATs propagate information across edge types and contexts, producing embeddings that capture both local interactions and broader biological structure.

Recent multimodal embedding methods (e.g., ProstT5 [6], CLASP [7]) highlight the value of integrating sequence, structure, and functional information, but they do not provide proteome-wide, context-specific embeddings for direct downstream use. Within this space, Pinnacle is one of the few publicly available methods producing pretrained, tissue- and cell-type–resolved embeddings [8]. Trained on a multiscale biological graph with a self-supervised objective, Pinnacle offers readily usable context-specific representations for tasks such as interaction prediction and target identification. However, its reliance on random data splits may overestimate generalization, particularly for understudied proteins [9].

A central challenge is that training and evaluation splits often fail to reflect real biological scenarios [10]. Because embeddings are learned directly from available evidence, data partitioning determines what information the model can access and what type of generalization it can achieve [11]. Random splits can inflate performance by allowing information leakage through network topology or shared annotations [12]. Moreover, biological networks are biased toward well-studied proteins [13], making it essential to evaluate models separately on well-studied and understudied subsets [14].

In this work, we introduce Gatsbi (Graph Attention with Split-Boosted Inference), a framework that integrates heterogeneous biological data into a unified network and learns context-aware embeddings using GAT-based message passing. We train under two biologically motivated data partitions—edge split and node split—and evaluate the resulting embeddings on interaction prediction, protein function prediction, and functional set prediction. We further assess performance across well-studied and understudied proteins. Across tasks, we find that performance depends strongly on the data split, and that biologically meaningful partitions reduce the performance gap between well-studied and understudied proteins compared with random splits. This work therefore addresses both the construction of protein embeddings and the evaluation conditions required for reliable assessment.

Key Takeaways:

1. Embedding construction and data partitioning strongly influence generalization. Varying the biological evidence available during training exposes how each split alters performance across tasks.
2. Separating well-studied and understudied proteins reveals how embeddings behave in data-limited regimes.
3. We provide pretrained embeddings for downstream prediction tasks.

## 2. Methods

### 2.1 Heterogeneous data sources capture complementary information across distinct functional aspects of protein behavior

#### 2.1.1 Protein sequence representations

We encode protein sequences using the Evolutionary Scale Modeling (ESM-2) protein language model, a large-scale transformer trained on millions of protein sequences to learn context-aware representations that capture evolutionary, structural, and functional properties of proteins [15]. For each protein, the corresponding amino acid sequence is passed through the pretrained ESM-2 model, and the resulting per-residue embeddings from the final hidden layer are aggregated using mean pooling to obtain a fixed-length protein-level representation. Thus, we obtain a vector representation for 20,074 human proteins. As these embeddings are derived from a pretrained model trained on large sequence corpora, they may encode information not controlled for by downstream evaluation splits, representing a potential source of pretraining-related bias.

#### 2.1.2 Human protein interaction network

The reference protein-protein interaction (PPI) network is the STRING network [16]. From the STRING database, we first obtain all human protein interactions and the associated score for each interaction. We use this data to obtain data on physical protein interactions, so we filter all STRING relationship to those that have experimental evidence of physical interaction. Reference PPI databases are known to have noisy interactions [17], and so we filter out interactions with an associated score of less than 0.6 [18]. The physical interaction subnetwork includes transferred edges (interologs inferred from orthologous species), though at our chosen score threshold these contribute minimally. This results in 16,462 number of protein nodes and 217,092 interactions.

#### 2.1.3 Protein co-expression patterns

Coordinated expression across biological contexts reflects shared regulatory control and pathway involvement, making co-expression a widely used indicator of functional coupling between proteins [19]. We obtain protein co-expression relationships from the STRING database using its co-expression evidence channel, which integrates large-scale transcriptomic and proteomic data across diverse biological contexts and converts these signals into calibrated functional association scores [16]. Again, we filter out relationships that have an association score of less than 0.6. This results in 11,205 proteins with 151,067 pairwise co-expression relationships.

#### 2.1.4 Tissue-specific functional associations

We additionally use HumanBase as a source of context-specific functional interaction networks [20]. HumanBase integrates bulk transcriptomic data with regulatory and interaction annotations to infer probabilistic gene–gene functional associations across 144 human tissue and cell-type contexts. We download the precomputed networks, retain high-confidence associations (posterior probability ≥ 0.6), and map gene identifiers to UniProt protein identifiers to obtain 17,617 proteins with 1,207,151 tissue-specific, functional relationships.

### 2.2 Integrating data sources into heterogeneous network

We model the complete set of pairwise protein relationships (interactions, expression, tissue-specific association) as a network, where the nodes correspond to proteins and an edge is present between any two nodes that have a pairwise relationship. We initialize the protein nodes with the sequence representation vector obtained from ESM-2. The edges are labeled with the data source from which the pairwise relationship originates (interaction, co-expression, tissue-specific association). If the edge originates from HumanBase, and thus is a tissue-specific relationship, the tissue type is also saved on the edge. Each edge is also annotated with the association score. The graph allows for multiple edges between the same pair of nodes, so a pair of proteins *u* and *v* can have multiple edges with different attributes (data source, weight, tissue type) connecting them. The final network has 18,049 nodes and 1,575,310 edges. Figure 1 displays a diagram describing the intergration of the data sources into one heterogeneous network.

**Figure 1.**
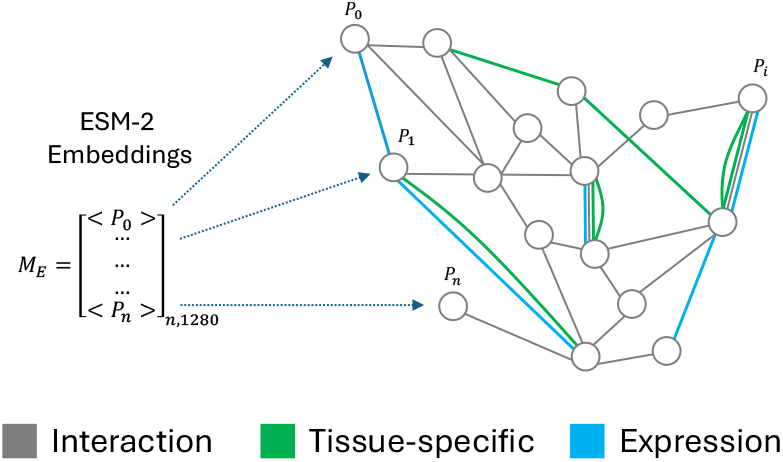
We constructed a heterogeneous protein network integrating sequence representations, physical interactions, co-expression, and tissue-specific associations. Nodes *P*_0_–*P*_*n*_ represent proteins in the human proteome and are initialized with ESM-2 embeddings *M*_*E*_ ∈ ℝ^*n*×1280^, providing one sequence vector per protein. Edges encode functional associations: gray edges denote physical interactions (STRING experimental channel), blue edges denote co-expression relationships (STRING co-expression channel), and green edges denote tissue-specific associations (HumanBase). Each edge is annotated with its data source, association score, and tissue type when applicable, and multiple labeled edges may connect the same protein pair.

### 2.3 Graph Attention Network

The constructed heterogeneous protein network is used as input to the Gatsbi graph attention network (GAT), which learns a low-dimensional embedding for each protein node by integrating the network information. The heterogeneous network is denoted as *G* = (*V, E*, 𝒯), where *V* is the set of protein nodes, *E* is the set of weighted edges, and 𝒯 is the set of edge types corresponding to physical interactions, co-expression, and tissue-specific functional associations. Each node *v* ∈ *V* is initialized with its ESM-2 sequence embedding 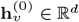.

At each GAT layer *𝓁*, node representations are updated via attention-based message passing over heterogeneous edge types. For a node *v*, the updated representation 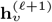 is computed as

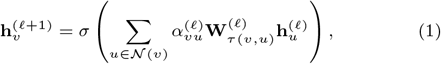

where *τ* (*v, u*) ∈ 𝒯 denotes the edge type between *v* and *u*, 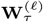 is a type-specific transformation matrix, and 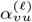 is the attention coefficient for message propagation along that edge. The attention coefficient is factorized as

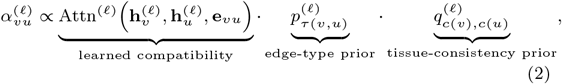

followed by normalization over the neighborhood of *v*.

Here, 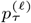 is a learnable scalar prior that controls the overall probability of traversing each edge type. The term 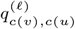 biases propagation toward edges whose tissue context matches the currently active tissue context *c*(*v*) of node *v*. When propagation follows a tissue-specific edge of type *t*, edges belonging to the same tissue context *t* are up-weighted relative to edges of other tissue contexts in subsequent message-passing steps, enforcing tissue-consistent information flow.

After *L* layers of propagation, the final embedding 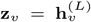 represents the learned protein embedding.

### 2.4 Training details for Gatsbi

Gatsbi learns protein representations from the heterogeneous network by optimizing a self-supervised link prediction objective that preserves relationships across data sources. During training, a subset of edges is masked, and the model is trained to predict their presence and type while discriminating against negative samples, using graphs that exclude the masked edges for training, validation, and testing. In the following subsections, we describe the construction of task-specific data splits and the downstream tasks used to evaluate the learned embeddings. Figure 2 A shows the model architecture overview, including the data split and the model.

**Figure 2.**
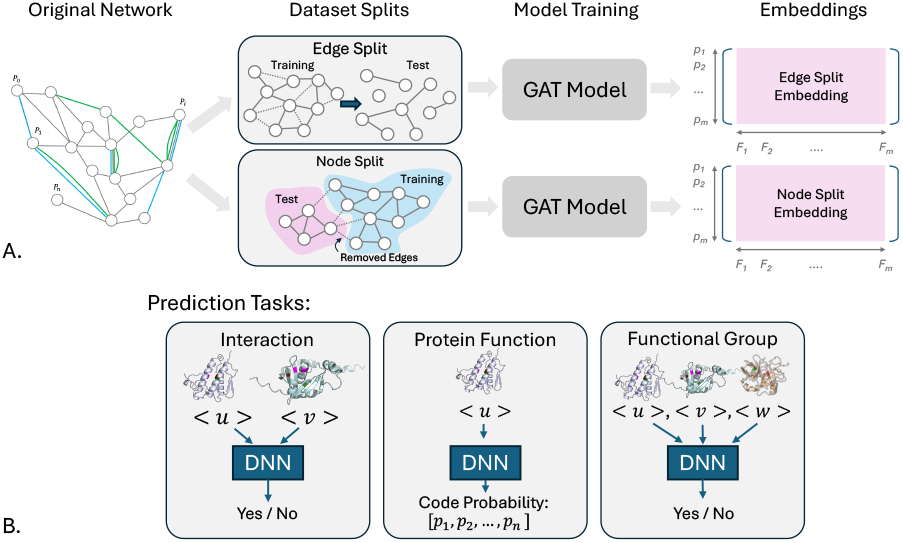
Overview of the evaluation framework. The original heterogeneous protein network is partitioned using either an edge split or a node split, producing distinct training and test graphs. A GAT model is trained on each split to generate protein embeddings, which are then used in downstream prediction tasks including interaction prediction, protein function classification, and functional group prediction.

#### 2.4.1 Task-driven graph splits for interaction-level and protein-level generalization

To rigorously evaluate generalization under different biological scenarios, we construct complementary data-splitting strategies - (1) edge (relationship-level) split and (2) node (protein-level) split. These correspond to established evaluation settings in protein interaction prediction (PMID: 23223166) [21], where the edge split aligns with the C1 setting (shared proteins across train and test), and the node split aligns with the C2 setting (proteins partitioned between train and test). We do not include a strict C3 setting, in which both proteins and their interactions are entirely absent from the training set, as this constitutes a distinct task involving prediction between proteins with no representation in the training graph and is not directly compatible with the embedding framework considered here.

##### Sequence homology control

To prevent information transfer through sequence similarity, we enforce a strict sequence identity threshold between training and test proteins. Pairwise sequence identity is computed using MMseqs2 [22], and proteins are partitioned such that no test protein exceeds 30% sequence identity with any training protein. This constraint is applied prior to constructing the node split and ensures that generalization cannot be achieved through trivial homology-based inference. This sequence identity constraint is applied only to the node split, where the goal is to evaluate generalization to entirely unseen proteins. In contrast, the edge split retains the full set of proteins by design, and evaluates recovery of unseen relationships among known proteins. Controlling for sequence homology in this setting is less critical, as proteins are already observed during training and homology-based transfer reflects realistic biological signal rather than information leakage.

##### Edge split (C1 setting)

In the edge split, we assign 70% of protein pairs to training and 30% to testing while retaining the full set of protein nodes and approximately preserving the per-datasource edge proportions. To minimize topological leakage, we construct the split such that, after removing all test edges, the shortest path between any test pair is at least 10 hops in the training graph. Because the network is a multigraph, we also group edges by protein pair: all relation types between a held-out pair are removed from training and placed exclusively in the test set.

##### Node split (inductive, C2 setting)

Proteins are partitioned into 70% training nodes and 30% test nodes. The constructed training graph contains only *V*_train_ and the edges among them, removing all edges incident to test nodes so that no interaction evidence about these proteins is available during representation learning. The link-prediction objective is optimized exclusively on this subgraph: a subset of training edges is masked, and the model is trained to recover them while discriminating against negative pairs sampled from *V*_train_, ensuring that no parameters depend on test-node information. To further prevent information transfer through sequence similarity, we enforce < 30% sequence identity between training and test proteins. At inference, the frozen encoder is applied inductively by inserting test proteins with their observed features and performing a forward pass using parameters learned from *V*_train_; link prediction involving test nodes is then performed without additional training. This setting evaluates generalization to genuinely understudied proteins that lack both interaction evidence and close sequence homologs during training, reflecting real-world scenarios in which new or sparsely characterized proteins must be interpreted using contextual structure learned from the rest of the network.

##### Training graph and loss definition

To unify the data-splitting strategies under a single training framework, we explicitly distinguish between: (1) the set of nodes available for message passing, *V*_mp_, (2) the set of edges available for message passing, *E*_mp_, and (3) the set of positive edges scored by the training loss, 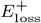.

For the edge split, all nodes participate in message passing, *V*_mp_ = *V*, while the message-passing graph contains only training edges, *E*_mp_ = *E*_train_; the same edges define the positive loss set, 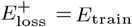. This configuration measures recovery of held-out relationships among known proteins.

For the node split (inductive), message passing is restricted to training proteins, *V*_mp_ = *V*_train_, with edges limited to those induced by these nodes, *E*_mp_ = *E*[*V*_train_]; the loss is likewise computed only on this subgraph, 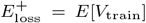. This configuration measures generalization to proteins absent during training.

##### Negative edge construction

For each training batch, negative edges are generated using a node-degree–matched sampling strategy rather than uniform random pairing. Specifically, for each positive edge in 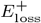, we sample non-adjacent protein pairs whose endpoint degrees closely match those of the corresponding positive pair, thereby controlling for degree-dependent biases in the link-prediction objective. Candidate negatives are drawn from *V*_mp_ while ensuring that they do not overlap with any known positive edges from the full network. We sample five such negatives per positive edge, maintaining a fixed negative-to-positive ratio of 5:1 across all experiments. In this setting, all pairs not observed as positive interactions are treated as candidate negatives, including both low-confidence and unobserved interactions.

#### 2.4.2 Gatsbi model configuration and training procedure

Using the data splits and optimization described above, we train the Gatsbi embedding model under a shared GAT-based link-prediction objective. For each split, the message-passing graph is constructed from the corresponding training portion of the network, using training edges for the edge split and the subgraph induced by *V*_train_ for the node split. Full model and training details are provided in Supplement Text 1.

### 2.5 Evaluating the protein embeddings

We evaluate the embeddings on three protein prediction tasks. Figure 2 B illustrates these three tasks.

#### 2.5.1 Prediction tasks used to measure the utility of protein embeddings

We evaluate the utility of the learned protein embeddings across three complementary prediction tasks. We run each experiment 10 times with random seeds. In all cases, we deliberately use lightweight predictive models so that performance reflects the quality of the embeddings rather than the capacity of the downstream classifier.

##### Interaction prediction (edge prediction)

To evaluate whether the embeddings encode information about physical and functional relationships, we perform protein–protein interaction prediction using BioGRID as the source of positive edges [23]. We filter all BioGRID edges to obtain a distinct set of edges from those included in the embedding creation. Each positive example corresponds to a curated physical interaction between two proteins. Negative examples are generated by randomly sampling protein pairs that do not appear in BioGRID. For each pair, we concatenate the two embedding vectors and train a shallow neural network classifier to predict whether an interaction exists. The task is evaluated under an edge-level split to ensure that test interactions involve protein pairs not seen during training.

##### Protein function prediction (node prediction)

We formulate enzyme function prediction as a multilabel node classification task using Enzyme Commission (EC) annotations obtained from UniProt [24]. Proteins with at least one EC label and an available embedding are retained. Each protein is represented by its learned embedding, and the task is evaluated under a node-level split to ensure that test proteins are unseen during training. A three-layer multilayer perceptron (MLP) is trained to predict the EC labels from the embedding vectors. This task assesses whether the embeddings capture functional biochemical properties of individual proteins.

##### Functional set prediction (pathway-level set classification)

To assess whether the embeddings capture higher-order biological organization, we evaluate prediction of functional protein sets derived from curated pathway annotations from Reactome [25]. Each positive example consists of the set of proteins assigned to a pathway. To construct negative examples, we generate “corrupted” pathways by replacing a fraction of proteins in a real pathway with randomly selected proteins, preserving pathway size while disrupting functional coherence. Because pathway sizes vary, we compute the median pathway size and pad or truncate each set to this fixed size. Protein embeddings within a set are aggregated using a multi-head attention pooling module, and the pooled representation is passed to a three-layer MLP to predict whether the set corresponds to a real pathway. The task is evaluated under a pathway-level split, ensuring that entire pathways in the test set are unseen during training.

Together, these tasks probe complementary aspects of the embedding space, providing a comprehensive evaluation of the biological information captured by the learned protein embeddings.

#### 2.5.2 Stratified Evaluation by Protein Evidence Level

To evaluate how well the learned embeddings support inference across different evidence regimes, we stratify proteins according to the amount of available biological knowledge. We quantify evidence using the total degree of each protein in the heterogeneous training network, which reflects both annotation density and interaction support. All 18,049 proteins are ranked by degree and partitioned into four equal-sized quantiles. The lowest-degree quantile (Q1; 4,512 proteins) represents understudied proteins with sparse connectivity and limited prior characterization, whereas the highest-degree quantile (Q4; 4,513 proteins) corresponds to well-studied proteins with extensive interaction and annotation evidence. For each prediction task, we report performance metrics separately for understudied and well-studied proteins. This stratified evaluation enables a direct assessment of how effectively the embeddings generalize to low-evidence proteins, a setting of particular practical importance given that many biologically and clinically relevant proteins remain poorly characterized.

To assess whether the degree-based proxy for studiedness aligns with external measures of protein characterization, we compared the Q1 and Q4 protein sets across several independent metrics, including Gene Ontology annotations [26], BioGRID interaction counts [23], Reactome pathway annotations [25], PubMed publication counts [27], and the Knownness/Unknome score [28]. Across these measures, proteins classified as well-studied by network degree consistently show higher annotation counts and knownness scores than those in the understudied group, indicating that the stratification broadly reflects biological characterization. A full quantitative comparison is provided in Supplement Table 2.

#### 2.5.3 Comparisons with baseline methods

##### Contextual embedding comparison

We benchmark Gatsbi against Pinnacle, a leading context-aware protein embedding method. Pinnacle is trained on a multiscale biological graph constructed from protein interaction networks derived from large-scale single-cell transcriptomic data, and optimized using a link-prediction objective. It produces context-specific embeddings for each protein across tissues and cell types, which are typically evaluated using random data splits.

The publicly available Pinnacle embeddings are used to derive a single per-protein representation by mean pooling across all contexts. These pooled embeddings are used directly in the same downstream pipelines as Gatsbi, ensuring that performance differences reflect the information encoded in the embeddings rather than task-specific retraining. The Pinnacle embeddings were obtained from the official implementation (GitHub: mims-harvard/PINNACLE, accessed Jan 2026).

To assess statistical significance, we perform a two-sided paired t-test across 10 runs for each model comparison, using a threshold of p < 0.05.

##### Task-specific model baselines

RAPPPID for interaction prediction [29], CLEAN for EC function prediction [30], and DeepSets for pathway-level set prediction [31]. These state-of-the-art models provide open-source implementations that can be retrained on our data splits and adapted to accept alternative input embeddings. For each method, we retain the original architecture and replace the input features with Gatsbi (edge- and node-split) or Pinnacle embeddings. CLEAN and DeepSets are used without modification beyond replacing the input embeddings, preserving their original training pipelines. Since RAPPPID’s Siamese AWD-LSTM operates on amino-acid sequences and cannot directly accept embedding inputs, we implement a RAPPPID-inspired variant: each protein embedding is projected into a shared latent space, the projected vectors are concatenated, and a classification head with Mish activations is applied.

## 3. Results

We report results for the experiments described in Sections 2.4 and 2.5, including performance of Gatsbi training under each data-splitting strategy and the effect of each split on all prediction tasks compared with baseline embedding methods.

### 3.1 Constructed heterogeneous graph captures context-specificity and adds more data for understudied proteins

Supplement Table 1 summarizes the number of nodes, edges, and connectivity statistics contributed by each individual data source as well as by the final heterogeneous graph. Figure 1 illustrates a representative subgraph of the heterogeneous network, highlighting the diverse edge types and context-specific relationships encoded between protein nodes. To understand how the distribution of edges in the understudied region, we plot a histogram of nodes at degree 0-100 in Figure 3. The heterogeneous graph contains fewer low-degree proteins than any single input network, indicating that proteins with limited experimental evidence in one data source are often supported by complementary evidence in others. As a result, the integrated graph offers improved context specificity and expanded information content for proteins that would otherwise remain poorly characterized. Figure 4 shows a histogram of the node degrees and the cutoffs of the well-studied (degree range 151-655) and understudied proteins (degree < 38).

**Figure 3.**
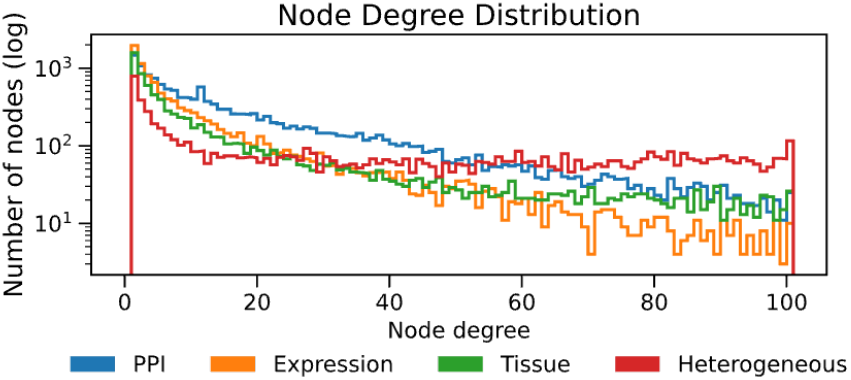
This figure shows a histogram of the edge degree for the nodes for each component data source and the combined heterogeneous graph. We see that the combined graph has the fewest low degree nodes, thus adding the most amount of information for the understudied proteins

**Figure 4.**
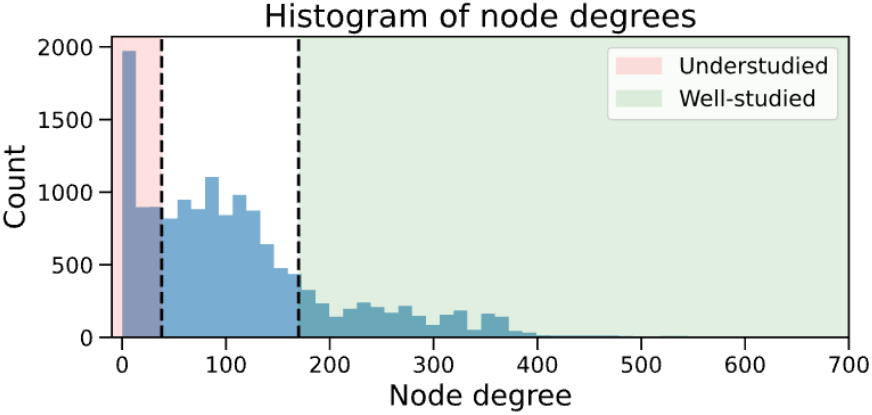
Proteins are grouped into well-studied and understudied categories based on node-degree thresholds (Section 2.5.2). The histogram shows the degree distribution with cutoffs marked by vertical lines. Understudied proteins (red) have degree ≤ 38, while well-studied proteins range from 151–655. In total, 3,705 proteins fall into the understudied group and 3,641 into the well-studied group.

### 3.2 Gatsbi embeddings generalize consistently

We evaluate the generalization performance of Gatsbi under the edge and node (inductive) data-splitting strategies described in Section 2.4.1.

Figure 5 shows t-SNE projections of the learned embedding space for the edge-split model. In Panel A, embeddings are colored by train, validation, and test membership; the lack of visible separation indicates that the model learns a unified representation space rather than partition-specific clusters. Panel B colors proteins by evidence level (well-studied in green, understudied in red, defined by degree quartiles in Section 2.5.2). Understudied proteins are broadly distributed and typically lie near well-studied proteins rather than forming isolated regions. Quantitatively, the average cosine distance between an understudied protein and its nearest well-studied neighbor is 0.234, compared to a random-background value of 0.225.

**Figure 5.**
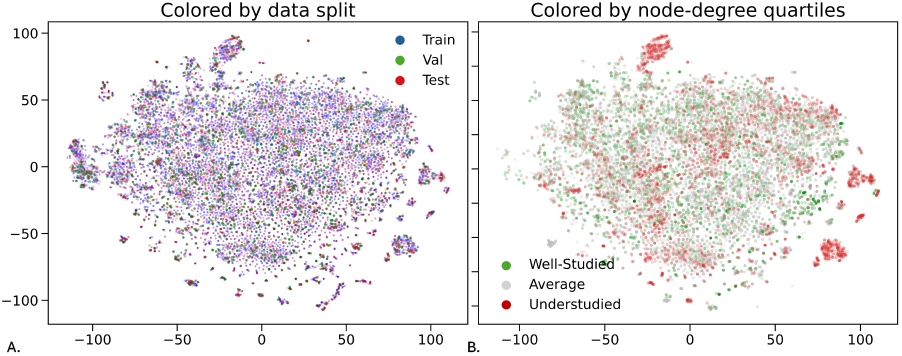
Embedding space learned by the GAT model (edge split). Embeddings colored by train, validation, and test sets show no partition-specific clustering, indicating good generalization. (B) Well-studied (green) and understudied (red) proteins, defined by node-degree quartiles, reveal a few understudied-enriched clusters, but most understudied proteins lie near well-studied neighbors. The average cosine distance to the nearest well-studied protein is 0.234, close to the random-baseline value of 0.225, indicating that understudied proteins occupy similar regions of the embedding space.

Supplement Figure 1 shows that the two distance distributions largely overlap, indicating that understudied proteins are not systematically farther from well-studied proteins in the embedding space.

Together, these results suggest that Gatsbi places low-evidence proteins near informative neighbors, supporting downstream annotation of understudied proteins through shared contextual structure.

### 3.3 Gatsbi embeddings improve downstream prediction across tasks and splits

#### Interaction prediction results

For interaction prediction, Gatsbi performed best under the edge split (AUROC 0.878, AUPRC 0.869; accuracy 0.794, F1 0.807) (Supplement Table 3), outperforming Pinnacle (AUROC 0.800, AUPRC 0.760). The inductive node split yielded lower scores (AUROC 0.746, AUPRC 0.730) but maintained high recall (0.853), indicating preserved sensitivity for unseen proteins. ROC and precision–recall curves (Figure 6) show the edge-split model consistently achieving stronger discrimination.

**Figure 6.**
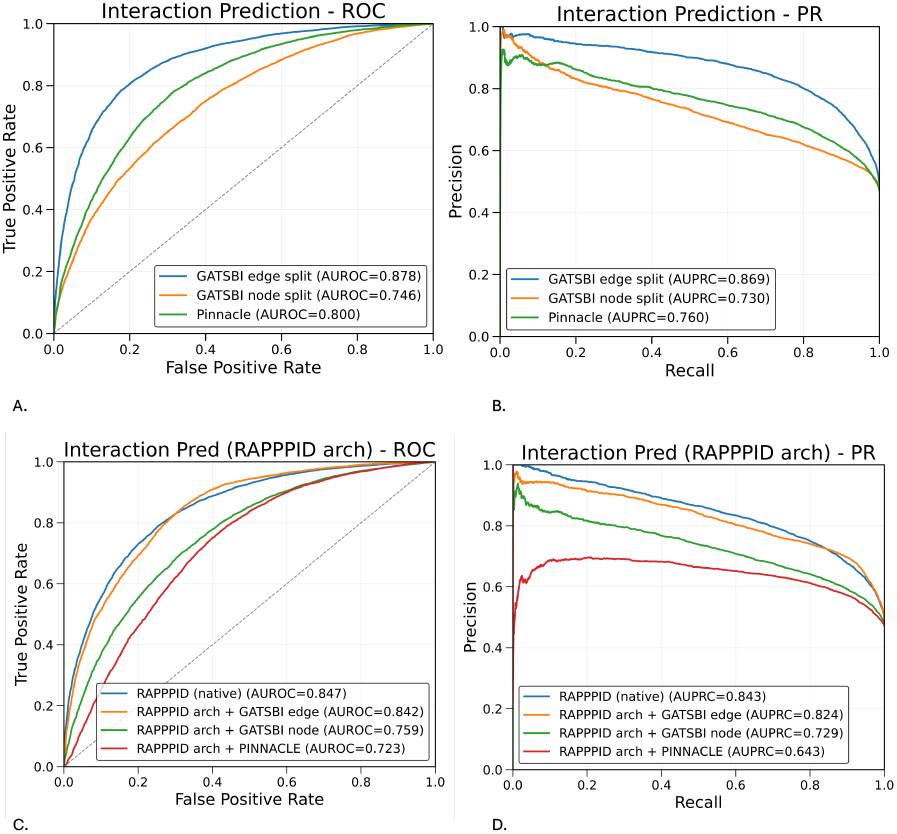
Interaction prediction performance. ROC (A) and precision–recall curves show that the Gatsbi edge-split model provides the strongest discrimination, reflecting its ability to capture interaction-relevant network structure. The node-split model remains competitive, indicating good generalization to unseen proteins. Architecture-swap results (C, D) show a similar pattern: supplying Gatsbi edge-split embeddings to the RAPPPID architecture yields performance close to RAPPPID’s native sequence-based model, whereas Gatsbi node-split and Pinnacle embeddings lead to larger drops, suggesting that the edge-split objective preserves pairwise interaction signals that transfer effectively to a dedicated interaction predictor.

#### Protein Function prediction results

For protein function prediction, Gatsbi achieved strong inductive performance (AUROC 0.679, AUPRC 0.231), comparable to Pinnacle (AUROC 0.668, AUPRC 0.197) (Supplement Table 4). The node-split model also showed higher accuracy (0.632 vs. 0.603) with similar micro-F1. The edge-split model performed worse (AUROC 0.592, AUPRC 0.175) despite high recall, reflecting reduced precision in the transductive regime. ROC and PR curves (Figure 7A–B) highlight these trends. Low AUPRC across methods reflects the extreme label imbalance of EC annotation, motivating micro-averaged evaluation. Overall, while Gatsbi offers only modest gains over Pinnacle, inductive training yields more informative representations for understudied proteins. Architecture-swap results (Figure 7C–D) show that the Clean model substantially outperforms all embedding-swap variants (AUROC 0.824, AUPRC 0.657). Replacing Clean’s learned features with Gatsbi node (0.687/0.304), Gatsbi edge (0.730/0.278), or Pinnacle embeddings (0.684/0.278) reduces performance, with the largest drop in AUPRC. This reflects Clean’s advantage in separating rare EC classes—a setting where a task-specific, contrastive objective benefits disproportionately from modeling extremely sparse labels.

**Figure 7.**
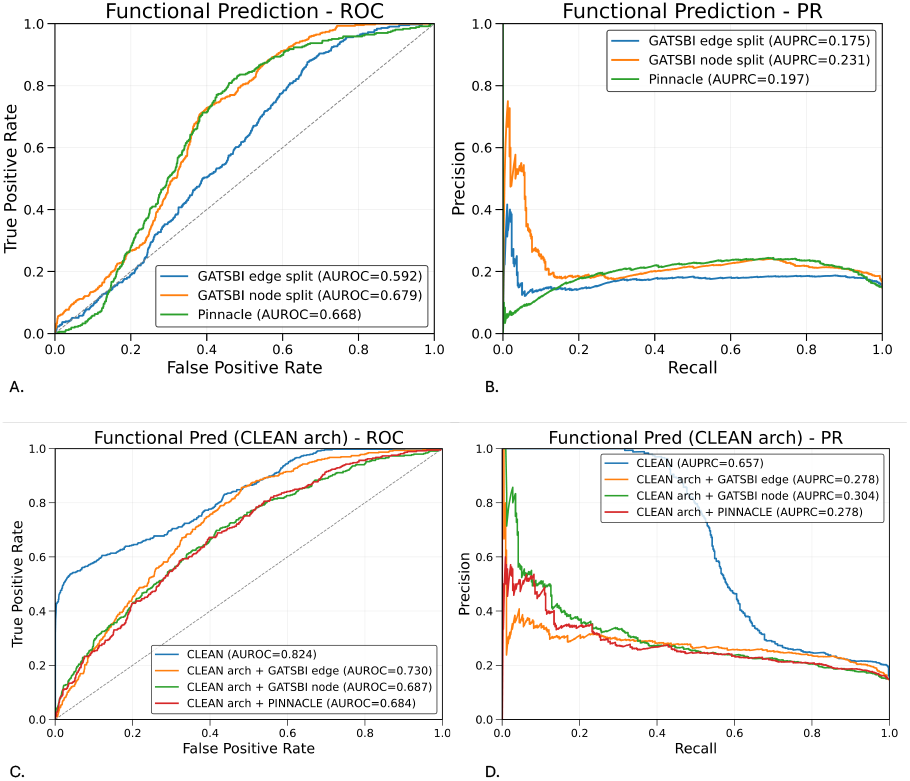
Protein function prediction. ROC (A) and precision–recall (B) curves show that the Gatsbi node-split model achieves stronger ranking performance than Pinnacle in both AUROC and AUPRC, despite uniformly low AUPRC values caused by extreme label imbalance. Architecture-swap results (C–D) show that the Clean model, which is specifically fine-tuned for this highly imbalanced multi-label EC task, substantially outperforms all embedding-swap variants, particularly in AUPRC. However, among the general-purpose embeddings, Gatsbi provides the strongest transfer performance

#### Functional Set prediction results

For functional set prediction, Gatsbi edge-split embeddings achieved the strongest performance (AUROC 0.804, AUPRC 0.821), outperforming Pinnacle (0.554/0.542) (Supplement Table 5). The node-split model performed lower (0.655/0.666) but remained clearly better than Pinnacle and retained high recall (0.897). ROC and PR curves (Figure 8A–B) show the edge-split model providing the most consistent discrimination. Architecture-swap results (Figure 8C–D) further support this: supplying Gatsbi edge embeddings to DeepSets improves performance over the native model (AUROC 0.746, AUPRC 0.756 vs. 0.644/0.634), indicating that graph-derived relational features transfer effectively to set-level prediction. Gatsbi node embeddings match DeepSets’ native performance, while Pinnacle underperforms. That Gatsbi edge embeddings surpass ESM within the same architecture suggests that interaction-aware graph training provides a stronger signal for identifying coherent protein sets than sequence-only features.

**Figure 8.**
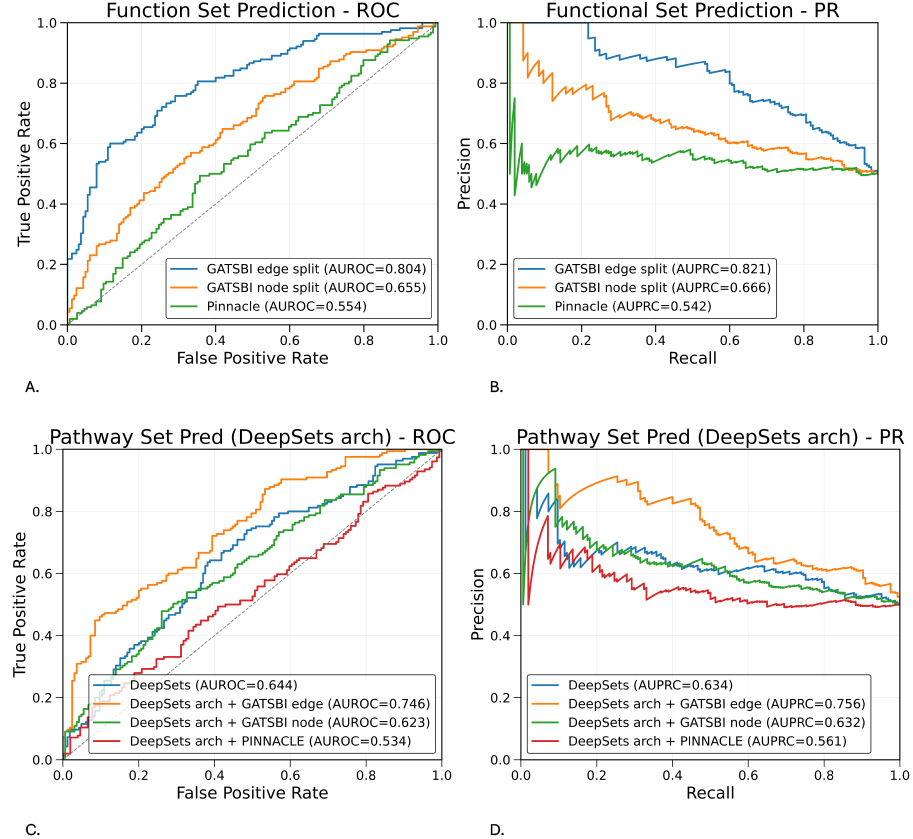
Functional set prediction. ROC (A) and precision–recall (B) curves show that Gatsbi edge-split embeddings provide the strongest discrimination of protein sets, capturing shared functional context more effectively than Pinnacle. Architecture-swap results (C–D) show that supplying Gatsbi edge embeddings to DeepSets improves performance over its native features, indicating that graph-derived relational information transfers well to set-level prediction and offers a stronger signal than sequence-only representations.

#### Ablation results

Ablations (Supplement Table 6) show that ESM-only embeddings provide a reasonable baseline but are consistently outperformed by graph-based variants. Incorporating network structure yields substantial gains, especially for interaction and set prediction, and combining interaction and expression graphs outperforms single-modality training. Random initialization degrades performance, confirming that pretrained sequence features are necessary but not sufficient. Together, these results show that Gatsbi’s improvements arise from integrating pretrained representations with structured, multi-modal biological networks.

### 3.4 Gatsbi performs higher than baseline method for understudied proteins

We further stratify evaluation results by protein degree quantiles, as described in Section 2.5.2. Performance stratified by protein evidence level confirms that Gatsbi provides greater benefit for understudied proteins than existing pretrained embeddings. Supplement Table 7) shows the performance metrics for each embedding method averaged across all three tasks. For understudied proteins, averaging across the three tasks, the edge-split model achieved AUROC 0.781 and AUPRC 0.801, exceeding Pinnacle (AUROC 0.522, AUPRC 0.511), while maintaining a balanced F1 of 0.724 compared with Pinnacle’s recall-driven behavior (F1 0.648, recall 0.928). The inductive node split also outperformed Pinnacle on ranking metrics (AUROC 0.632 vs. 0.522; AUPRC 0.641 vs. 0.511), demonstrating that G atsbi embeddings transfer more effectively to proteins lacking training evidence. Similar trends were observed for well-studied proteins, where the edge split reached AUROC 0.822 and AUPRC 0.836, again surpassing Pinnacle (AUROC 0.578, AUPRC 0.566). Quantitatively, the improvement over Pinnacle is largest in the understudied regime (ΔAUROC = +0.259; ΔAUPRC = +0.290) but remains substantial for well-studied proteins (ΔAUROC = +0.244; ΔAUPRC = +0.270), indicating consistent gains across evidence levels. Precision–recall behavior further highlights this difference: Pinnacle attains high recall at the expense of precision, whereas Gatsbi achieves a more balanced operating point with higher F1. These results are consistent with the embedding-space analysis, where understudied proteins are positioned near informative well-studied neighbors, enabling effective knowledge transfer. The stratified performance per task is visualized in Figure 9 A. The performance deltas (Figure 9 B) show that Gatsbi provides the largest gains over Pinnacle for understudied proteins in interaction and functional set prediction, highlighting improved transfer to proteins with limited prior evidence. In contrast, for function prediction, the improvement is similar across evidence levels, reflecting the inherently harder multiclass task where label imbalance, rather than protein coverage, constrains performance.

**Figure 9.**
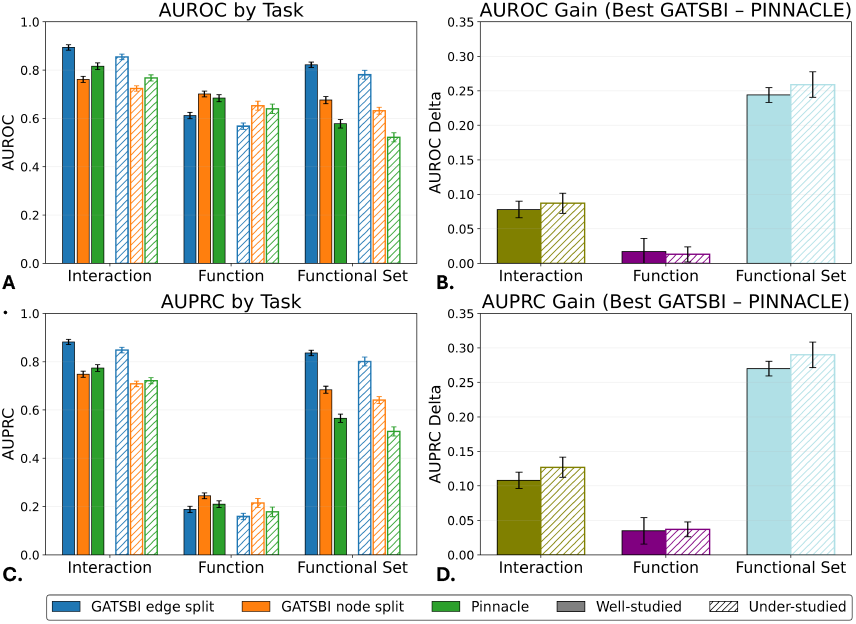
Performance of Gatsbi and Pinnacle for well-studied and understudied proteins across three tasks. Solid bars denote well-studied proteins; hatched bars denote understudied proteins. (A) Stratified AUROC for interaction, function, and functional-set prediction shows that Gatsbi outperforms Pinnacle in both groups, with larger gains for understudied proteins. (B) AUROC deltas (best Gatsbi – Pinnacle) highlight the largest improvements for understudied proteins in interaction and functional-set prediction. (C) Stratified AUPRC similarly favors Gatsbi, especially for understudied proteins. (D) AUPRC deltas show the same pattern, with the biggest gains for understudied proteins in interaction and functional-set prediction, while function prediction remains constrained by multilabel imbalance.

We further investigated the biological relevance of false positive predictions, particularly focusing on high-confidence errors, as these may reflect missing or incomplete annotations rather than true model failures. Figure 10 highlights two such predictions. Panel A shows a predicted interaction between the understudied proteins Protocadherin-15, involved in cochlear function, and Stereocilin, required for hair-cell stereocilia formation. Although this interaction is not documented in humans, a related association has been reported in mole-rats [32], suggesting a conserved but underreported functional relationship. Panel B shows a predicted functional set in which amyloid beta precursor protein (APP) was added to a module containing neuroserpin, PDCD10, alpha-synuclein, and major prion proteins. These proteins share roles in neuronal development and cell-cycle regulation, and APP is a neuronal surface receptor that promotes synaptogenesis. While the original network lacked a direct APP–neuroserpin link, an association has been described by Yepes *et al*. [33]. Together, these examples illustrate that many false positives are enriched for shared biological pathways, cellular context, or functional coherence, indicating that they may correspond to plausible but currently unannotated relationships rather than true errors.

**Figure 10.**
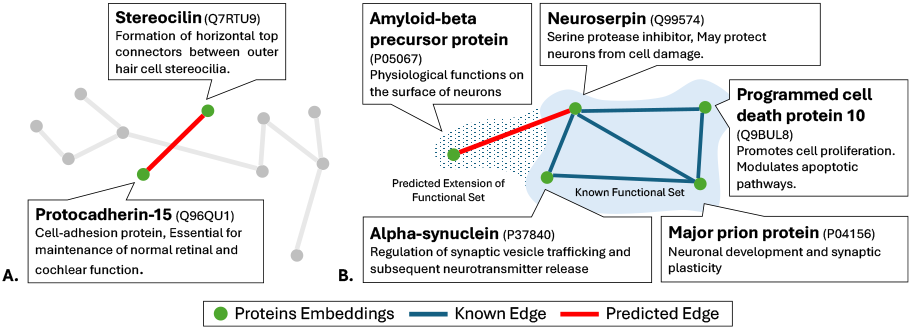
Visualization of two example model predictions. Green nodes denote proteins, red edges are model-predicted relationships, and blue edges are known associations. These examples illustrate how apparent false positives can reflect biologically plausible but previously unreported links, especially for understudied proteins. (A) Predicted interaction between Protocadherin-15 and Stereocilin, both involved in cochlear hair-cell function. (B) Predicted functional set in which APP is added to a module containing neuroserpin, PDCD10, *α*-synuclein, and prion proteins, all associated with neuronal development and cell-cycle regulation.

## 4. Discussion

This study shows that how protein embeddings are trained and evaluated strongly shapes their apparent utility for biological prediction. By integrating heterogeneous evidence—interactions, co-expression, tissue context, and sequence features—into a unified graph attention framework, Gatsbi learns representations that outperform existing pretrained embeddings across interaction, function, and functional set prediction. More importantly, the results highlight that evaluation protocol—not just model design—substantially alters the conclusions one might draw about embedding quality.

Evaluations restricted to well-studied proteins overestimate real-world performance. Pinnacle introduced the first large-scale context-aware protein embeddings and demonstrated the value of heterogeneous biological context. Gatsbi builds on this foundation with training and evaluation strategies aligned to practical use cases. When stratified by evidence level, Gatsbi shows clearer advantages for understudied proteins, achieving a more balanced precision–recall profile than Pinnacle. Embedding-space analyses support this behavior: understudied proteins cluster near informative neighbors, enabling more effective transfer of contextual information. The contrast between edge and node splits further underscores that different biological questions require different evaluation regimes. The edge split measures recovery of missing relationships among known proteins and favors tasks like interaction and set prediction. The inductive node split instead reflects scenarios where new proteins must be annotated without prior interaction evidence. The reversal of performance trends across these splits shows that no single benchmark captures all relevant use cases. For function prediction, improvements were more modest, reflecting the difficulty of extreme multilabel imbalance. The persistent gap between AUROC and AUPRC suggests that future work should incorporate label-aware decoders or hierarchical modeling rather than relying solely on generic classifiers.

Several limitations remain. Although the node split reduces leakage through topology and sequence similarity, other dependencies—such as shared experimental biases—are harder to control. Our framework also relies on pretrained ESM-2 embeddings, which may encode information not fully isolated by downstream splits. Addressing this would require retraining language models under controlled data partitions. Pooling Pinnacle’s context-specific embeddings into a single vector may underrepresent its strengths in context-dependent settings. Finally, our evaluation focuses on human proteins; extending to additional species would provide a broader test of cross-organism generalization.

Overall, these results show that progress in protein representation learning depends as much on evaluation design as on model architecture. By distinguishing transductive and inductive regimes and stratifying by evidence level, Gatsbi provides a more realistic assessment of practical utility—particularly for the long tail of understudied proteins that remain central to functional genomics.

## Data availability

Code and models for our experiments are available at https://github.com/Helix-Research-Lab/GATSBI-embedding

## Author contributions statement

GN and RBA discussed the experiments. GN conducteed the experiments, analyzes the results, and wrote the manuscript. GN and RBA editted the manuscript.

## References

1. R. Morris, K. A. Black, and E. J. Stollar, “Uncovering protein function: from classification to complexes,” vol. 66, pp. 255–285.

2. X. Wang, D. Bo, C. Shi et al., “A survey on heterogeneous graph embedding: Methods, techniques, applications and sources,” vol. 9, pp. 415–436.

3. C. Tran, S. Khadkikar, and A. Porollo, “Survey of protein sequence embedding models,” vol. 24, p. 3775.

4. G. Nayar and R. B. Altman, “Heterogeneous network approaches to protein pathway prediction,” vol. 23, pp. 2727–2739.

5. A. G. Vrahatis, K. Lazaros, and S. Kotsiantis, “Graph attention networks: A comprehensive review of methods and applications,” vol. 16, p. 318.

6. M. Heinzinger, K. Weissenow, J. G. Sanchez et al., “Bilingual language model for protein sequence and structure,” vol. 6, p. lqae150.

7. N. Bolouri, J. Szymborski, and A. Emad, “Multi-modal protein representation learning with CLASP,” p. 2025.08.10.669533.

8. M. M. Li, Y. Huang, M. Sumathipala et al., “Contextual AI models for single-cell protein biology,” vol. 21, pp. 1546– 1557.

9. J. Bernett, D. B. Blumenthal, D. G. Grimm et al., “Guiding questions to avoid data leakage in biological machine learning applications,” vol. 21, pp. 1444–1453.

10. M. Milano, G. Agapito, and M. Cannataro, “Challenges and limitations of biological network analysis,” vol. 11, p. 24.

11. E. Ursu, A. Minnegalieva, P. Rawat et al., “Training data composition determines machine learning generalization and biological rule discovery,” vol. 7, pp. 1206–1219.

12. Y. Shao, H. Li, X. Gu et al., “Distributed graph neural network training: A survey,” vol. 56, pp. 1–39.

13. T. G. Brooks, N. F. Lahens, A. Mrčela et al., “Challenges and best practices in omics benchmarking,” vol. 25, pp. 326– 339.

14. N. Bordin, C. Dallago, M. Heinzinger et al., “Novel machine learning approaches revolutionize protein knowledge,” vol. 48, pp. 345–359.

15. Z. Lin, H. Akin, R. Rao et al., “Evolutionary-scale prediction of atomic-level protein structure with a language model,” vol. 379, pp. 1123–1130.

16. D. Szklarczyk, K. Nastou, M. Koutrouli et al., “The STRING database in 2025: protein networks with directionality of regulation,” vol. 53, pp. D730–D737.

17. T. Rout, A. Mohapatra, and M. Kar, “A systematic review of graph-based explorations of PPI networks: methods, resources, and best practices,” vol. 13, p. 29.

18. A. Nogueira-Rodríguez, D. Glez-Peña, C. P. Vieira et al., PPI prediction from sequences via transfer learning on balanced but yet biased datasets: An open problem, ser. Lecture Notes in Networks and Systems. Springer Nature Switzerland, pp. 31–40.

19. J. D. Montenegro, “Gene co-expression network analysis,” vol. 2443, pp. 387–404.

20. R. S. G. Sealfon, C. L. Theesfeld, J. Funk et al., “HumanBase: an interactive AI platform for human biology,” pp. 1–2.

21. Y. Park and E. M. Marcotte, “Flaws in evaluation schemes for pair-input computational predictions,” vol. 9, pp. 1134– 1136.

22. M. Steinegger and J. Söding, “MMseqs2 enables sensitive protein sequence searching for the analysis of massive data sets,” vol. 35, pp. 1026–1028.

23. R. Oughtred, J. Rust, C. Chang et al., “The BioGRID database: A comprehensive biomedical resource of curated protein, genetic, and chemical interactions,” vol. 30, pp. 187–200.

24. A. G. McDonald and K. F. Tipton, “Enzyme nomenclature and classification: the state of the art,” vol. 290, pp. 2214– 2231.

25. M. Milacic, D. Beavers, P. Conley et al., “The reactome pathway knowledgebase 2024,” vol. 52, pp. D672–D678.

26. M. Ashburner, C. A. Ball, J. A. Blake et al., “Gene ontology: tool for the unification of biology. the gene ontology consortium,” vol. 25, pp. 25–29.

27. M. Knapp, “PubMed features to save your time,” vol. 24, pp. 1–9.

28. J. J. Rocha, S. A. Jayaram, T. J. Stevens et al., “Functional unknomics: Systematic screening of conserved genes of unknown function,” vol. 21, p. e3002222.

29. J. Szymborski and A. Emad, “RAPPPID: towards generalizable protein interaction prediction with AWD-LSTM twin networks,” vol. 38, pp. 3958–3967.

30. T. Yu, H. Cui, J. C. Li et al., “Enzyme function prediction using contrastive learning,” vol. 379, pp. 1358–1363.

31. M. Zaheer, S. Kottur, S. Ravanbakhsh et al., “Deep sets.”

32. S. J. Pyott, M. van Tuinen, L. A. Screven et al., “Functional, morphological, and evolutionary characterization of hearing in subterranean, eusocial african mole-rats,” vol. 30, pp. 4329–4341.e4.

33. M. Yepes, Y. Woo, and C. Martin-Jimenez, “Plasminogen activators in neurovascular and neurodegenerative disorders,” vol. 22, p. 4380.

