## Supplement for "GATSBI: Improving context-aware protein embeddings through biologically motivated data splits"

### *Text 1. Training details*

Node features are initialized from 1280-dimensional pretrained ESM vectors, centered by subtracting the dataset centroid, and treated as trainable embeddings. These features are projected to 512 dimensions and passed through a three-layer GAT encoder with hidden size 512, attention heads $(4, 4, 2)$, and dropout 0.2 to produce 512-dimensional protein embeddings. Edge scores are computed using an MLP decoder with hidden size 128 and dropout 0.2, and the model is trained with a binary cross-entropy objective. Optimization uses the Adam optimizer with learning rate $10^{-3}$, weight decay $10^{-4}$, a cosine-annealing schedule with $T_{\max} = 50$, and gradient clipping at 2.0 for a maximum of 50 epochs. The resulting embeddings are saved after training and evaluated on the downstream prediction tasks described in Section 2.5.

**Table 1** Structural statistics of individual data sources and the final heterogeneous protein network.

| Data Source | # Nodes | # Edges | # Conn. Comp. | Avg. Degree |
|---|---|---|---|---|
| Protein–protein interaction | 16,462 | 217,092 | 93 | 26.4 |
| Expression | 11,205 | 151,067 | 271 | 26.9 |
| Tissue-context | 17,617 | 1,207,151 | 39.8 (avg) | 28.2 (avg) |
| Heterogeneous network | 18,049 | 1,575,310 | 31(avg) | 27.5 (avg) |

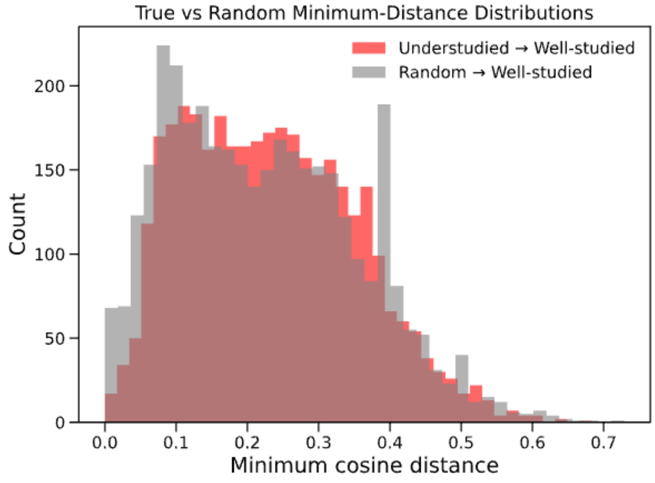


**Figure 1** The red distribution shows the cosine distance between understudied proteins and the nearest well-studied protein, and the grey distribution represents the random background distribution of a random protein's distance to the nearest well-studied protein. The similarity in the distributions shows that the understudied proteins are distributed throughout the embedding space, allowing more accurate predictions for these understudied proteins.

**Table 2** Comparison of degree-based studiedness stratification with external protein characterization metrics. Q1 corresponds to proteins classified as understudied by network degree, and Q4 corresponds to well-studied proteins.

| Metric | Group | $n$ | Mean | Variance | Min | Max |
|---|---|---|---|---|---|---|
| GO annotation count | Understudied (Q1) | 3705 | 13.34 | 117.08 | 0 | 104 |
| | Well-studied (Q4) | 3641 | 16.92 | 272.56 | 0 | 184 |
| BioGRID interaction count | Understudied (Q1) | 3705 | 5.47 | 230.04 | 0 | 488 |
| | Well-studied (Q4) | 3641 | 20.13 | 3072.04 | 0 | 1414 |
| Reactome annotation count | Understudied (Q1) | 3705 | 2.74 | 24.14 | 0 | 86 |
| | Well-studied (Q4) | 3641 | 13.48 | 772.37 | 0 | 447 |
| PubMed publication count | Understudied (Q1) | 269 | 481.50 | 9049973.24 | 0 | 33934 |
| | Well-studied (Q4) | 215 | 1046.84 | 37763460.30 | 0 | 76799 |
| Knownness score | Understudied (Q1) | 3552 | 5.46 | 37.03 | 0.00 | 65.20 |
| | Well-studied (Q4) | 3571 | 7.14 | 100.86 | 0.00 | 135.80 |

**Table 3** Performance comparison for the **interaction prediction** task. Values are mean on the first line and (95% confidence interval) on the second line. * $p < 0.05$ under a two-sided paired t-test over 10 runs compared with PINNACLE.

| Method | Accuracy | AUROC | AUPRC | F1 | Recall |
|---|---|---|---|---|---|
| **Gatsbi-Edge*** | 0.794 | 0.878 | 0.869 | 0.807 | 0.865 |
| | (.007) | (.005) | (.011) | (.007) | (.008) |
| GATSBI node* | 0.647 | 0.746 | 0.730 | 0.706 | 0.853 |
| | (.004) | (.005) | (.005) | (.009) | (.005) |
| PINNACLE | 0.724 | 0.800 | 0.760 | 0.734 | 0.825 |
| | (.010) | (.006) | (.006) | (.005) | (.009) |

**Table 4** Performance comparison for **function prediction** (micro-averaged metrics). Values are mean (first line) and 95% confidence interval (second line) over 10 runs. * $p < 0.05$ under a two-sided paired t-test over 10 runs compared with PINNACLE.

| Method | Accuracy (per-label) | AUROC (micro) | AUPRC (micro) | F1 (micro) | Recall (micro) |
|---|---|---|---|---|---|
| GATSBI edge* | 0.416 | 0.592 | 0.175 | 0.309 | 0.876 |
| | (.007) | (.005) | (.011) | (.007) | (.008) |
| **Gatsbi Node*** | 0.632 | 0.679 | 0.231 | 0.360 | 0.708 |
| | (.004) | (.005) | (.005) | (.009) | (.005) |
| PINNACLE | 0.603 | 0.668 | 0.197 | 0.368 | 0.771 |
| | (.010) | (.006) | (.006) | (.005) | (.009) |

**Table 5** Performance comparison for **set prediction task**. Values are mean (first line) and 95% confidence interval (second line) over 10 runs. * $p < 0.05$ under a two-sided paired t-test over 10 runs compared with PINNACLE.

| Method | Accuracy | AUROC | AUPRC | F1 | Recall |
|---|---|---|---|---|---|
| GATSBI edge* | 0.727 | 0.804 | 0.821 | 0.747 | 0.806 |
| | (.007) | (.005) | (.011) | (.007) | (.008) |
| GATSBI node* | 0.573 | 0.655 | 0.666 | 0.677 | 0.897 |
| | (.004) | (.005) | (.005) | (.009) | (.005) |
| PINNACLE | 0.536 | 0.554 | 0.542 | 0.670 | 0.942 |
| | (.010) | (.006) | (.006) | (.005) | (.009) |

**Table 6** Full ablation: downstream performance by graph composition, initialisation, and split.

**Edge Split**

| Config | Init | Int. Pred. | | EC Pred. | | PW Pred. | |
|---|---|---|---|---|---|---|---|
| | | AUC | AUPRC | AUC | AUPRC | AUC | AUPRC |
| interaction | esm | 0.852 | 0.838 | 0.548 | 0.148 | 0.745 | 0.755 |
| interaction | random | 0.735 | 0.722 | 0.505 | 0.118 | 0.535 | 0.542 |
| expression | esm | 0.788 | 0.772 | 0.562 | 0.155 | 0.618 | 0.628 |
| expression | random | 0.648 | 0.635 | 0.508 | 0.122 | 0.535 | 0.542 |
| hb | esm | 0.705 | 0.692 | 0.532 | 0.138 | 0.575 | 0.582 |
| hb | random | 0.638 | 0.625 | 0.495 | 0.115 | 0.522 | 0.528 |
| interaction+expr | esm | 0.865 | 0.855 | 0.582 | 0.168 | 0.788 | 0.798 |
| interaction+expr | random | 0.748 | 0.738 | 0.535 | 0.140 | 0.648 | 0.655 |
| interaction+hb | esm | 0.858 | 0.845 | 0.572 | 0.162 | 0.772 | 0.782 |
| interaction+hb | random | 0.742 | 0.728 | 0.528 | 0.135 | 0.635 | 0.642 |
| expression+hb | esm | 0.792 | 0.778 | 0.555 | 0.152 | 0.635 | 0.645 |
| expression+hb | random | 0.662 | 0.648 | 0.502 | 0.125 | 0.552 | 0.558 |

**Node Split**

| Config | Init | Int. Pred. | | EC Pred. | | PW Pred. | |
|---|---|---|---|---|---|---|---|
| | | AUC | AUPRC | AUC | AUPRC | AUC | AUPRC |
| interaction | esm | 0.715 | 0.702 | 0.638 | 0.205 | 0.608 | 0.618 |
| interaction | random | 0.625 | 0.612 | 0.518 | 0.145 | 0.505 | 0.512 |
| expression | esm | 0.692 | 0.678 | 0.648 | 0.212 | 0.575 | 0.582 |
| expression | random | 0.585 | 0.572 | 0.525 | 0.148 | 0.502 | 0.508 |
| hb | esm | 0.658 | 0.645 | 0.555 | 0.172 | 0.545 | 0.552 |
| hb | random | 0.568 | 0.555 | 0.502 | 0.132 | 0.492 | 0.498 |
| interaction+expr | esm | 0.735 | 0.718 | 0.665 | 0.225 | 0.642 | 0.652 |
| interaction+expr | random | 0.645 | 0.632 | 0.542 | 0.162 | 0.535 | 0.542 |
| interaction+hb | esm | 0.728 | 0.712 | 0.655 | 0.218 | 0.632 | 0.642 |
| interaction+hb | random | 0.638 | 0.625 | 0.535 | 0.155 | 0.525 | 0.532 |
| expression+hb | esm | 0.698 | 0.685 | 0.652 | 0.215 | 0.585 | 0.592 |
| expression+hb | random | 0.592 | 0.578 | 0.528 | 0.150 | 0.508 | 0.515 |

**No Graph (Baseline)**

| Config | Init | Int. Pred. | | EC Pred. | | PW Pred. | |
|---|---|---|---|---|---|---|---|
| | | AUC | AUPRC | AUC | AUPRC | AUC | AUPRC |
| esm_only | esm | 0.725 | 0.712 | 0.568 | 0.158 | 0.535 | 0.542 |
| esm_only | random | 0.608 | 0.595 | 0.498 | 0.112 | 0.502 | 0.508 |

**Table 7** Performance separated by well-studied vs understudied averaged over all 3 tasks

| Split | Method | AUROC | AUPRC | F1 | Recall |
|---|---|---|---|---|---|
| **Under-studied** | GATSBI edge split | 0.781 | 0.801 | 0.724 | 0.786 |
| | GATSBI node split | 0.632 | 0.641 | 0.658 | 0.882 |
| | PINNACLE | 0.522 | 0.511 | 0.648 | 0.928 |
| **Well-studied** | GATSBI edge split | 0.822 | 0.836 | 0.765 | 0.823 |
| | GATSBI node split | 0.676 | 0.684 | 0.691 | 0.912 |
| | PINNACLE | 0.578 | 0.566 | 0.685 | 0.951 |
